# cytometree: a binary tree algorithm for automatic gating in cytometry analysis

**DOI:** 10.1101/335554

**Authors:** Daniel Commenges, Chariff Alkhassim, Raphael Gottardo, Boris Hejblum, Rodolphe Thiébaut

## Abstract

**Motivation:** Flow cytometry is a powerful technology that allows the high-throughput quantification of dozens of surface and intracellular proteins at the single-cell level. It has become the most widely used technology for immunophenotyping of cells over the past three decades. Due to the increasing complexity of cytometry experiments (more cells and more markers), traditional manual flow cytometry data analysis has become untenable due to its subjectivity and time-consuming nature.

**Results:** We present a new unsupervised algorithm called `“cytometree”` to perform automated population discovery (aka gating) in flow cytometry. `cytometree` is based on the construction of a binary tree, the nodes of which are subpopulations of cells. At each node, the marker distributions are modeled by mixtures of normal distribution. Node splitting is done according to a normalized difference of Akaike information criteria (AIC) between the two models. Post-processing of the tree structure and derived populations allows us to complete the annotation of the derived populations. The algorithm is shown to perform better than the state-of-the-art unsupervised algorithms previously proposed on panels introduced by the Flow Cytometry: Critical Assessment of Population Identification Methods (FlowCAP I) project. The algorithm is also applied to a T-cell panel proposed by the Human Immunology Project Consortium (HIPC) program; it also outperforms the best unsupervised open-source available algorithm while requiring the shortest computation time.

**Availability:** An R package named “cytometree” is available on the CRAN repository.

**Supplementary information:** Supplementary data are available.

## 1 Introduction

Recent technological advancements in instrumentation have transformed the field of flow cytometry by enabling rapid, multidimensional quantification of millions of individual cells to define cellular subpopulations and assess cellular heterogeneity (O’Neill *et al*., 2013; Aghaeepour *et al*., 2013). Traditionally, flow cytometry data are analyzed manually by drawing geometric shapes (referred to as ‘gates’) around populations of interest in a series of 1-2 dimensional data representations. This process, known as gating, is time-consuming and highly subjective (Aghaeepour *et al*., 2013). Modern instruments including both flow and mass cytometers are now capable to quantify between 20 and 50 proteins, leading to high-dimensional data that are impossible to exhaustively explore by manual analysis. Several supervised and unsupervised algorithms have been proposed for automatic gating, including model-based clustering approaches (Lo *et al*., 2008; Finak *et al*., 2009; Chan *et al*., 2008; Pyne *et al*., 2009; Qian *et al*., 2010; Aghaeepour *et al*., 2011; Cron *et al*., 2013a), a nonparametric density-based approach (Naumann and Wand, 2009), and a spectral clustering algorithm (Zare *et al*., 2009), among others. A number of these have been compared in the open competition set-up by the FlowCAP consortium (Aghaeepour *et al*., 2013) and many developments have followed as a result (Brinkman *et al*., 2015, 2016; Finak *et al*., 2014) as reviewed in Saeys *et al*. (2016). Automated cell classification has also been applied to mass cytometry by time of flight (CyTOF) data (Lee *et al*., 2017). Many of these algorithms performed rather well on the FlowCAP benchmark data. However, no single method was uniformly superior on all datasets. Additionally, some of these methods were very computationally demanding and no method led to biologically interpretable cell populations (i.e. population labels are exchangeable). To overcome these problems, supervised algorithms including flowDensity (Malek *et al*., 2015) and OpenCyto (Finak *et al*., 2014) have been proposed and compared to manual gating for several panels of cells analyzed by nine laboratories by the Human Immuno Phenotyping Consortium (HIPC). However, being supervised, these approaches require significant tuning and restrict the exploration of flow cytometry data to pre-specified cell populations.

Here, we propose a new method that is fast, compares favorably to state-of-the-art approaches, and leads to biologically interpretable populations. It uses the same basic idea that experimentalists utilize when analyzing data: a given cell either expresses or does not express a given protein (i.e. the marginal distribution of each marker is mostly bimodal). That is, for most markers, the cells will be either negative (-) or positive (+). We approximate the distribution of each marker by a mixture of two normal distributions. This process allows us to cycle through all markers, and to build a binary tree, the leaves of which are the terminal subpopulations. The annotation is completed using a post-processing procedure. We call this new method `“cytometree”`.

Our paper is organized as follows: We first present the `cytometree` algorithm, then an illustration of the outputs of the program using the HIPC T-cell panel. Finally, we demonstrate its utility and performance on FlowCAP I and FlowCAP III challenge benchmark data.

## 2 Methods

### 2.1 Principle

The distribution of each marker is modeled as a mixture of two normal distributions. A criterion, noted *D*, is computed for each marker to measure the improvement brought by the mixture over a single normal. The marker with the highest value of the criterion is chosen to be the first node of the tree. The population of cells is divided into two subsets, provisionally annotated negative (-) and positive (+) for this marker. The same procedure is then applied for each subpopulation, and thus a binary tree is obtained. The tree growth is stopped when the highest value of the criterion is below a pre-specified threshold. The leaves of the tree are the final subpopulations obtained by this algorithm. Branches of the tree (i.e. the gating path) lead to a biologically interpretable annotation (e.g. CD3+/CD4+) for each subpopulation given the markers that were used at each node in the path (from the root to the given leaf). However, a given path may not make use of all available markers, as some markers might not exhibit bimodality, or have a *D* value always lower than other markers in competition. As such, we also propose a post-processing annotation algorithm to generate alternative population labels that make use of all markers.

### 2.2 Binary tree algorithm based on difference of normalized AIC

In this section, we describe more precisely the construction of the binary tree. Let us denote by *s*_*j*_ the set of available markers at node *j*, that is, the set of markers that have not been used for defining parent nodes of *j*, and by *n*_*j*_ the number of cells at node *j*. At a given node *j*, `cytometree` inspects whether the population of cells can be split by exhaustively searching over all available markers in the set *s*_*j*_, the marker for which the fluorescence distribution of the observed cells maximizes the criterion *D*, defined as

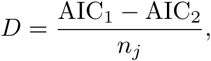

where AIC_1_ and AIC_2_ are the conventional Akaike criterion values for the one and two component mixture models, respectively. *D* is a normalized version of the difference of Akaike criterion (AIC) (Commenges *et al*., 2008). The advantage of this criterion is that it does not depend on *n*_*j*_, but estimates the difference of Kullback-Leibler divergences from the true distribution (the true distribution is the distribution from which the data are supposed to be generated). A difference of 0.1 was considered “large”. Thus, for each marker *m ∈ s*_*j*_, the distribution of the fluorescence intensity 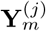 can be modeled as either a single normal with mean *μ* and variance *σ*^2^ or a two component mixture model as follows:

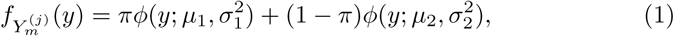

where *ϕ*(.; *μ, σ*^2^) is the normal density of mean *μ* and variance *σ*^2^. The parameters of the mixture 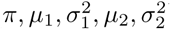 are estimated by maximum likelihood using an EM algorithm (Fraley and Raftery, 2002; Fraley *et al*., 2012). Then, the criterion for marker *m*, *D*_*m*_, is computed using the likelihoods of the one and two component mixture models. The maximum value of *D* over all markers is defined as *D*^*^ = max_*m*_ *D*_*m*_. If *D*^*^ is above a pre-specified threshold *t*^*^, the population at node *j* is split according to the values of the marker *m*^*^ that achieved this maximum and two child nodes are obtained. Cells with 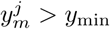 (resp. 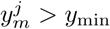) form the subpopulation of the left (resp. right) child, where 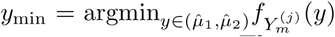. If *D*\* < *t*\* the tree growth is stopped, and *j* is a leaf of the tree. The threshold *t*^*^ can be adjusted to find more or fewer populations; our experience showed that values between 0.1 and 0.2 generally give good results.

When the tree has stopped growing, the leaves yield a partition of the data into *L* subpopulations 𝒫= {*S*_1_,…, *S*_*L*_..

It can be shown that the algorithm runs in linearithmic fashion as a function of the number of cells *n*, that is, the complexity is in 𝒪 (*n* log *n*) (see Figure S6 in Supplemental material for an empirical check). The computational cost increases linearly with the dimension (number of markers) for each node. Moreover, the computational cost is linear in the number of nodes and the number of nodes is lower than twice the number of leaves, that is the sub-populations. The number of leaves of the tree is likely to increase slightly with number of dimensions but the number of sub-populations cannot be very high. Assuming the number of sub-population is bounded, the complexity is essentially linear in the dimension.

### 2.3 Annotation algorithm

Given that the binary tree construction is unsupervised and depends on a pre-specified threshold *t*^*^, some of the available markers may not have been used to find the different cell subpopulations (i.e. some markers in some paths never pass the threshold, or always have *D* values lower than other markers). To recover a complete annotation using all available markers, we devised a post-processing exhaustive annotation method. This step can be supervised or unsupervised. In the supervised option, the number of expression levels of each marker is fixed by the user, in the unsupervised option it is proposed by the algorithm based on the *D* criterion.

*Supervised option*. In the supervised option, we wish to annotate the subpopulations for some or all of the markers. In general, we wish to annotate the subpopulations as positive or negative for the chosen markers. For each chosen marker, we rank the means of the *L* subpopulations, forming (*S*_(1),_…, *S*_(*L*)_). Then we form *L –* 1 partitions of the *L* subpopulations into two groups: the partition *p*, for *p* = 1,…, *L –* 1, is 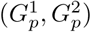 with 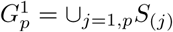 and 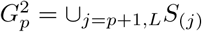 Then we find the best partition *G*_*p*⋆_ in the sense of minimizing the within-cluster variance:

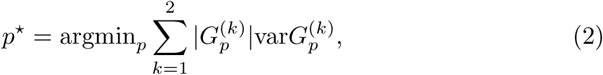

where 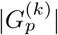 is the cardinal of 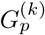. This is the same formula that is used in the K-means algorithm, but only *L –* 1 partitions are tried and the observations are one-dimensional (since we work marker-by-marker). Thus, this step of the algorithm is very fast. Finally, we label subpopulations (*S*_(1)_,…, *S*_(*p*⋆)_) as negative “(–)” and (*S*_(*p*⋆)_),…, *S*_(*L*)_) as positive “(+)” for the marker at hand.

We can perform the same type of algorithm for partitioning the subpopulations into three groups, “-”, “+” and “++” for some markers. Here the number of partitions is (*L–*1)(*L–*2)/2. This can also be done for the markers used in the tree. As an example, we may wish to find three levels of CD45RA; this is exemplified in HIPC Patient 12828 replicate 3 from the NHLBI dataset; see Supplementary Material S1.

*Unsupervised option*. In this option, for the markers not used in the tree, we compute the *D* criterion comparing the fits of the marginal distribution obtained by one normal distribution and by a mixture of two or three normal distributions for judging whether there are one, two, or three groups. For the markers used in the tree, we compute the *D* criterion to compare the fits obtained by a mixture of two and three normal distributions.

### 2.4 *F*-measure

The *F* -measure is a popular metric to evaluate clustering methods. It can be used as a way to summarize the concordance between two classification methods (one being set as the reference). This measure is the harmonic mean of precision and recall (Aghaeepour *et al*., 2013). The precision is the number of cells correctly assigned to a given cluster divided by the total number of cells assigned to this cluster. The recall is the number of cells correctly assigned to a given cluster divided by the number of cells that should be assigned to this cluster according to the reference method. The total *F* -measure is then calculated for each combination of the reference clusters and the predicted clusters. It yields a value of [0, 1], with 1 indicating a perfect match between the two clustering methods.

### 2.5 Benchmarking

#### 2.5.1 FlowCAP I challenge

Several unsupervised algorithms have been compared to manual gating done by a consensus of 8 manual operators (from 8 different laboratories) on 5 data sets. These data sets included four human data sets: graft-versus-host disease (GvHD), diffuse large B-cell lymphoma (DLBCL), symptomatic West Nile virus (WNV), and normal donors (ND); the fifth was a mouse data set (hematopoietic stem cell transplant (HSCT)). Each of the 5 data sets includes multiple samples of up to 10^5^ cells measured on a maximum of 10 markers. The results were set to be used as benchmark data in the FlowCAP I challenge (Aghaeepour *et al*., 2013). The data were downloaded from the FlowCAP project website as part of the FlowCAP I challenge.

#### 2.5.2 HIPC T-cell panel

The Human Immune Phenotyping Consortium (HIPC) was developed with the aim of standardizing flow cytometry immunophenotyping in clinical studies. Finak *et al*. (2016) investigated whether automated gating could help standardizing flow cytometry data analysis. We used a part of the data collected in this study to assess the performance of the `cytometree` algorithm, focusing on the T-cell panel. Seven laboratories (or centers) stained three replicates of three cryopreserved PBMC samples and returned usable FCS files to the main center for manual and automated gating. The automated gating used a combination of algorithms including flowDensity, which is a supervised algorithm. Data sets are publicly available from the ImmuneSpace database (Brusic *et al*., 2014) and were used as part of the FlowCAP III challenge.

We reproduced the variability analysis of the estimated proportions *p*_*rij*_ in replicate *r* of sample *i* in center *j*, of each subpopulation of cells presented in Finak *et al*. (2016). Denoting by *Y*_*rij*_ = log *p*_*rij*_*/*(1– *p*_*rij*_), the logit of these proportions, the model was as follows:

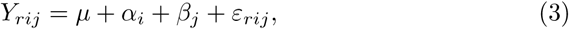

where *μ* is the intercept, *α*_*i*_ is a sample random effect, *β*_*j*_ is a center random effect, and *ε*_*rij*_ is the residual error. All these random variables are assumed to be independent normal with zero means and with variances 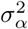, 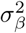 and 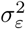, respectively. This model allows us to estimate and quantify the different sources of variability. There is one difficulty in this analysis in that the logit of zero is infinite. For this reason, as in Finak *et al*. (2016), we excluded zero values.

## 3 Results

### 3.1 Example of a T-cell sample analysis

For the purpose of illustrating how the algorithm works and what the output looks like, we show some results for a single T-cell sample: the Stanford FCS data for sample 1349, replicate 3 from the HIPC dataset (Finak *et al*., 2016). The fits for the single normal and the mixture of two normals are computed for all markers, and the differences of normalized AIC (*D* values) are computed. Figure 1 shows the fits obtained with the mixture compared to non-parametric fits obtained by a kernel method; note the very good fit obtained for the CD4 by the mixture of two normal distributions. The CD4 had the best *D* criterion (1.31); the first node is declared to be CD4 and is labeled “CD4.1”. Cells are then separated into provisionally negative or positive CD4 groups. Again, the values of the *D* criterion are computed for all markers except CD4 in the two populations; in both cases, CCR7 wins. Thus, two nodes are created, “CCR7.2” and “CCR7.3”; the fits of the mixture are shown for these two distributions. The tree growth continues until the maximum *D* criterion value is smaller than 0.1. The tree obtained is displayed in Figure 2.

**Figure 1:**
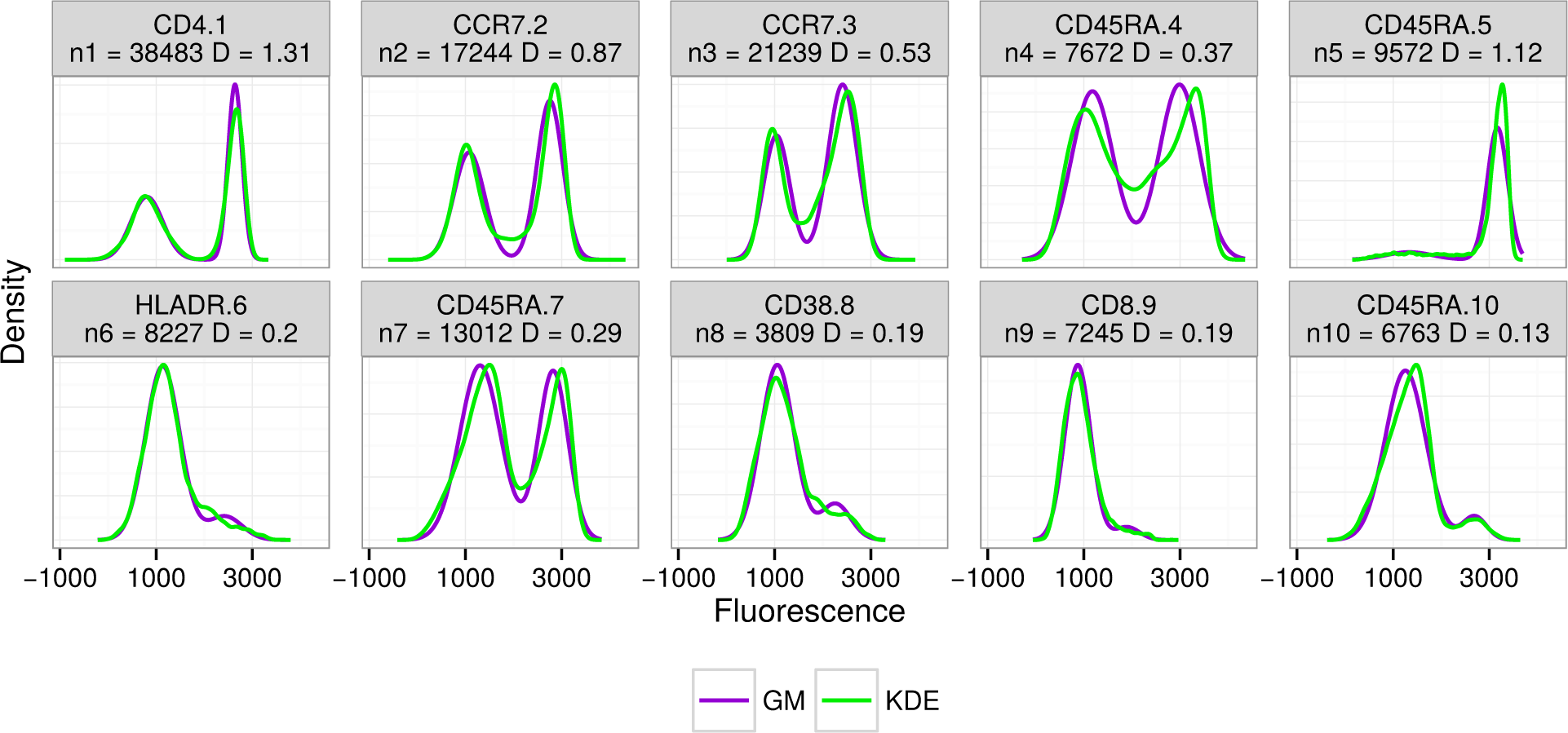
Conditional marginal node distributions for the T cells of patient 1349 in the Stanford FCS data set, replicate 3. In blue, fits obtained with mixtures, in green, non-parametric fits obtained by a kernel method.

**Figure 2:**
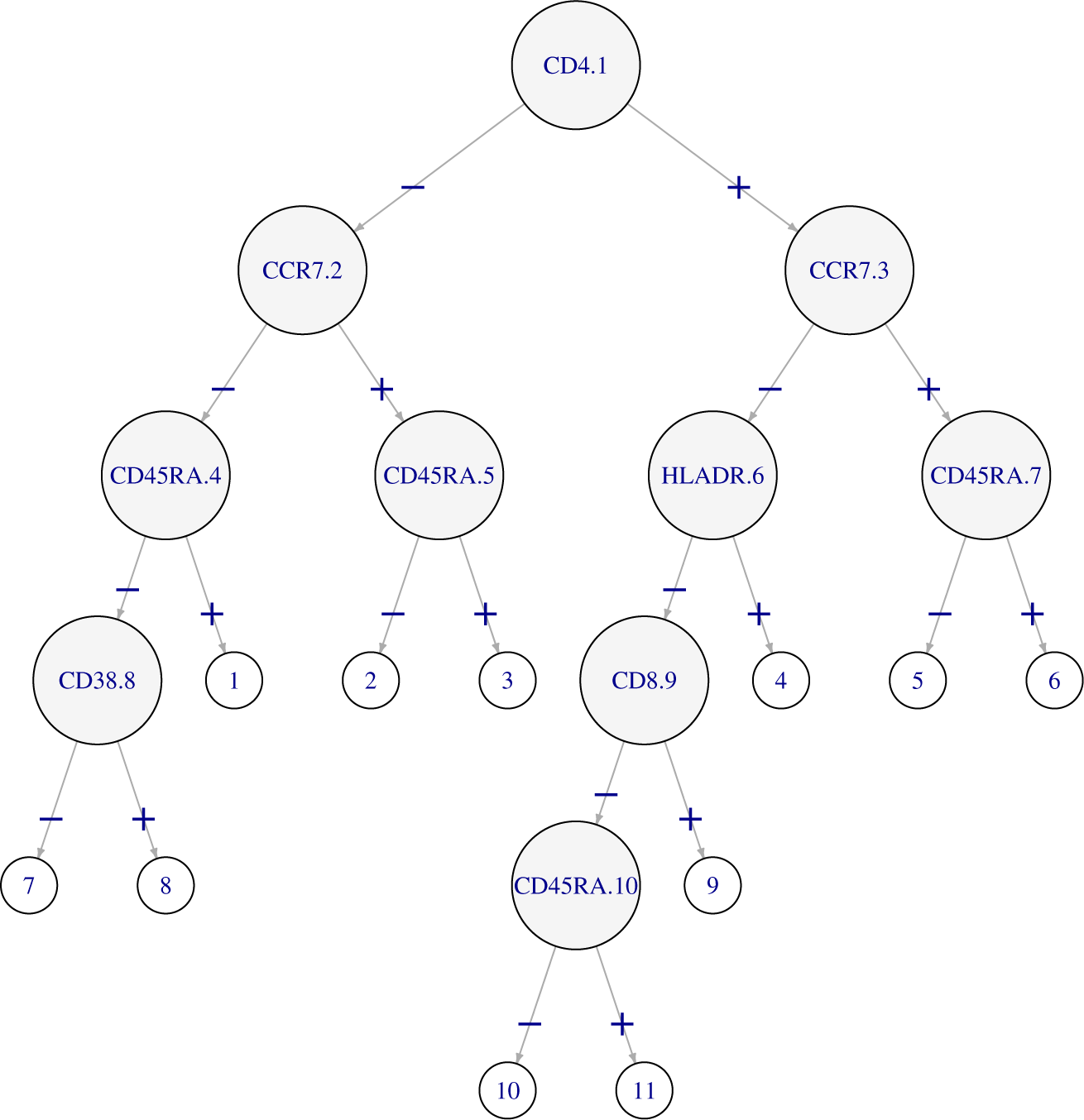
Partitioning tree for the T cells of patient 1349 (replicate 3) from the Stanford dataset. Each node that has children is labeled with the marker on which the subpopulation is split; leaves are numbered.

Although `cytometree` approximates univariate distributions by mixtures of normal distributions, it is possible to reconstitute a bivariate scatter plot such as that analyzed visually in manual gating. The scatter plots obtained for CD45RA and CCR7 for CD4+ cells by manual gating and `cytometree` are displayed in Figure 3, which shows that the two are almost identical for two patients; however for patient 12828a, `cytometree` fails to split the CCR7 population in CD45RA+ and CD45RA-due to the very small number of cells that may constitute this subpopulation.

**Figure 3:**
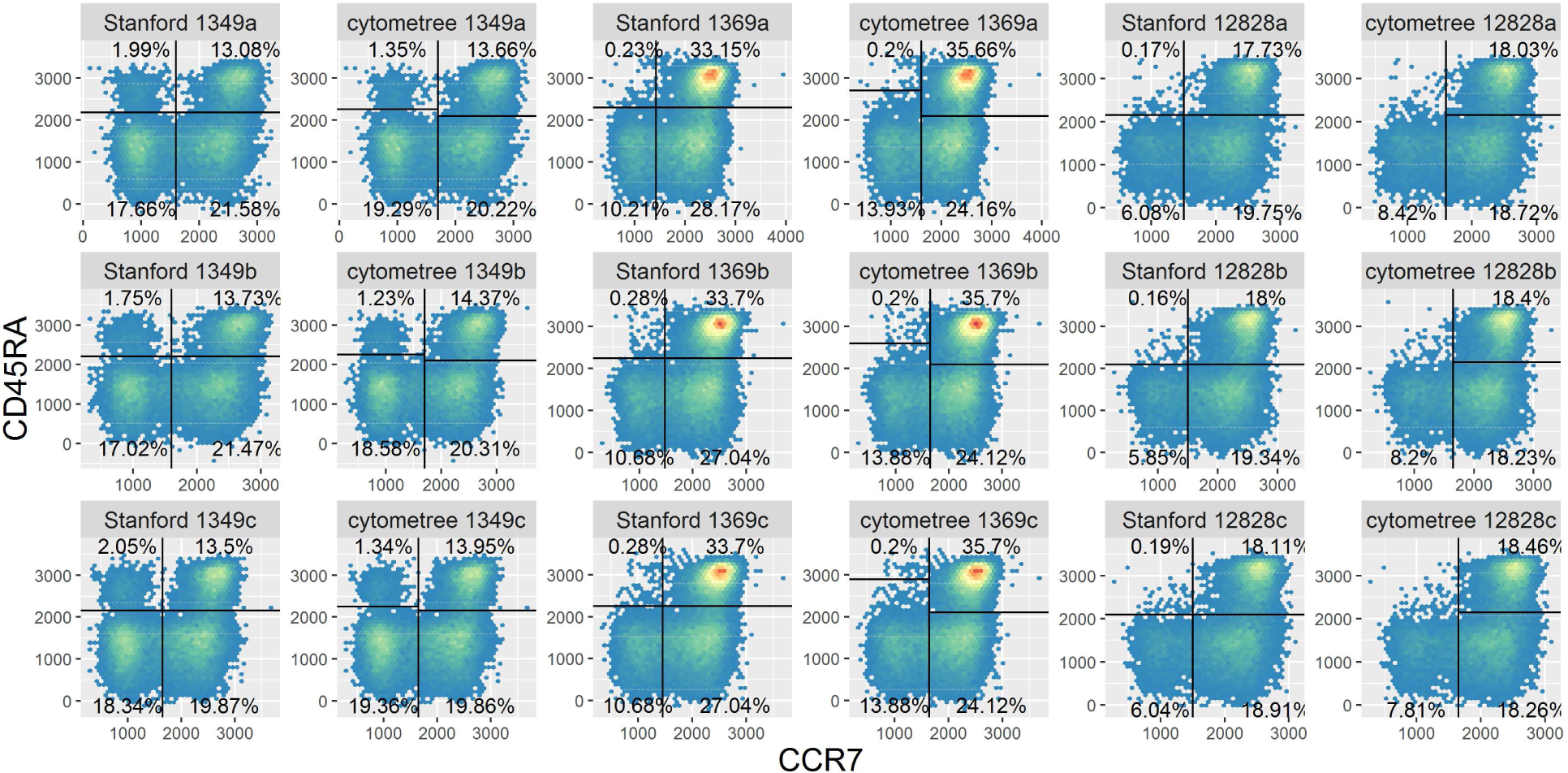
Plots of hexagonally binned data showing four subpopulations of the T-CD4 cells (top left: Effector T cells, top right: Naive T cells:, bottom left: Effector memory T cells, bottom right: Central memory T cells). Gating from Stanford is compared to that obtained by cytometree for the three patients and their three replicates.

The next step of the algorithm is the annotation process. This is necessary because although the binary tree gives an annotation for each explored sub-population, the annotation remains incomplete as some markers may be left unused in the tree growth process. So for all markers, we apply the annotation algorithm described in Section 2.3 to gather the found subpopulations in two or three groups. Results for patient 1349, replicate 3 are displayed in Figure 4, which shows the distribution of the markers for all the subpopulations found, and the result of the clustering algorithm. The subpopulations found in the tree are thus annotated, and a table is constructed to describe them and give the proportion of each subpopulation in the sample. For the chosen sample, the results are shown in Table 1, together with the proportions that were found for the same sample by manual gating in the Stanford center.

**Table 1:** T-cell subpopulations evaluated by Stanford for patient 1349 (replicate 3): proportions estimated by manual gating and by `cytometree` are given.

| **Population Name** | **Corresponding Markers** | **cytometree** | **Stanford** |
| --- | --- | --- | --- |
| CD4 Activated | CD3+ CD8- CD4+ CD38+ HLADR+ | 2.55% | 1.39% |
| CD8 Activated | CD3+ CD8+ CD4- CD38+ HLADR+ | 1.54% | 1.67% |
| CD4 Central Memory | CD3+ CD8- CD4+ CCR7+ CD45RA- | 19.86% | 19.87% |
| CD8 Central Memory | CD3+ CD8+ CD4- CCR7+ CD45RA- | 2.85% | 4% |
| CD4 Effector | CD3+ CD8- CD4+ CCR7- CD45RA+ | 1.34% | 2.05% |
| CD8 Effector | CD3+ CD8+ CD4- CCR7- CD45RA+ | 10.04% | 7.94% |
| CD4 Effector Memory | CD3+ CD8- CD4+ CCR7- CD45RA- | 16.23% | 18.34% |
| CD8 Effector Memory | CD3+ CD8+ CD4- CCR7- CD45RA- | 8.36% | 9.85% |
| CD4 Naive | CD3+ CD8- CD4+ CCR7+ CD45RA+ | 13.95% | 13.5% |
| CD8 Naive | CD3+ CD8+ CD4- CCR7+ CD45RA+ | 22.03% | 21.39% |

**Figure 4:**
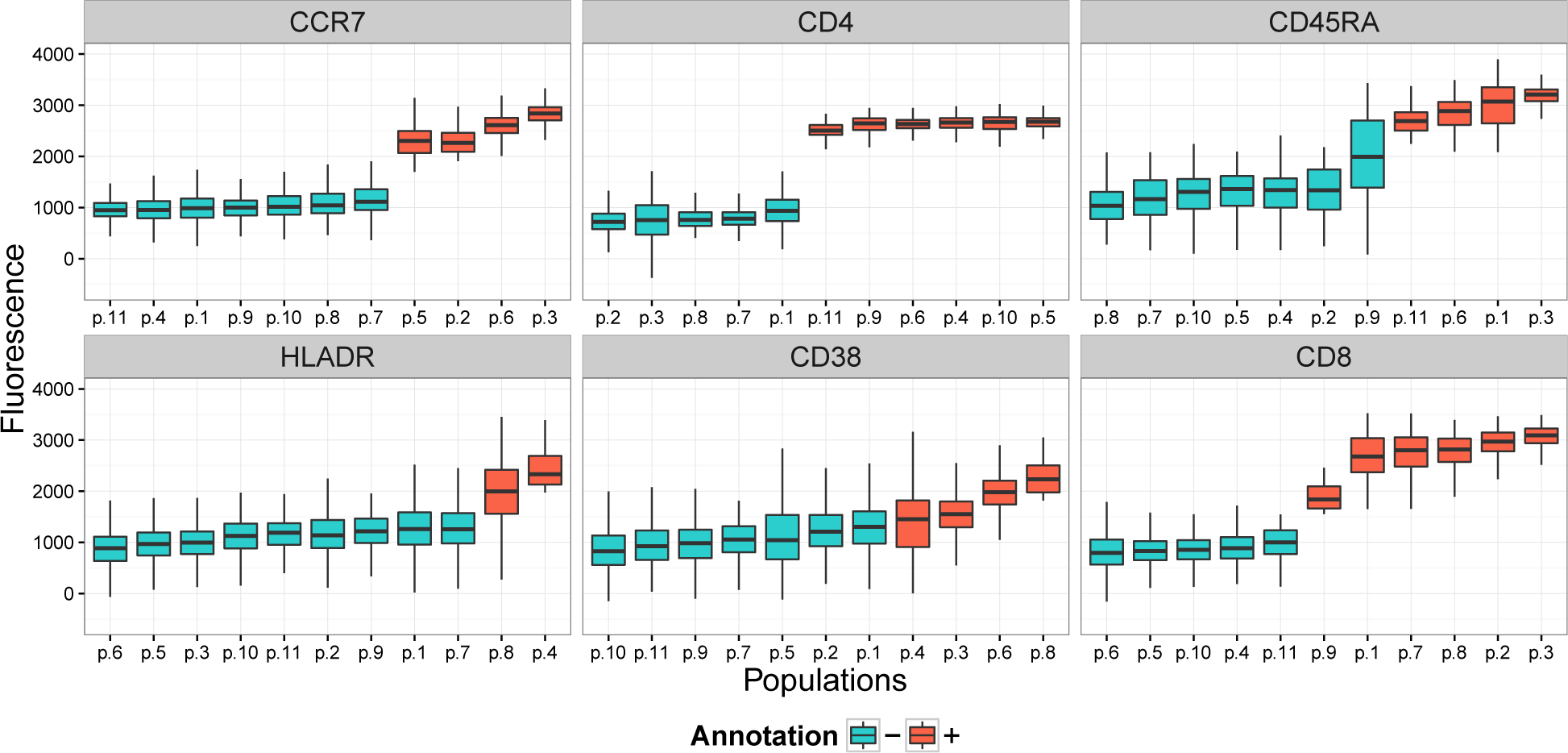
Example of annotation results for the T cells of patient 1349 (replicate c) from the Stanford dataset. Abscissas show the populations from the underlying tree. The clustering algorithm allows allocating the populations into “-” and “+” groups for each marker.

### 3.2 cytometree obtains the best results in FlowCAP I

Table 2 shows the performance of `cytometree` on the FlowCAP I data, compared to the four best performing methods reviewed by Aghaeepour *et al*. (2013): ADICyt (Chan *et al*., 2008), flowMeans (Aghaeepour *et al*., 2011), FLOCK (Qian *et al*., 2010) and FLAME (Pyne *et al*., 2009). The *F* -measures were computed for all samples available for a given dataset and the mean over all samples is reported. The best open source unsupervised algorithm in the FlowCAP I study appears to be flowMeans. `cytometree` nearly always obtains the highest values, and the mean *F* -measure is 0.90 for the default value *t*^*^ = 0.1, making it the best unsupervised approach in the completely automated challenge. We explored a range of values of *t*^*\**^ between 0.05 and 0.25; the F-measure is rather stable on this range. In terms of computing time `cytometree` was one of the fastest algorithms, even faster than flowMeans, taking on average about one minute per sample.

**Table 2:** F-measures for cytometree, with t* respectively equal to ^1^0.05, ^2^0.1, ^3^0.15, ^4^0.2, ^5^0.25 and the four algorithms that performed the best on the FlowCAP I challenge data sets. Mean F-measures and mean run times are also given.

| Method | GvHD<br>(n=12) | HSCT<br>(n=30) | DLBCL<br>(n=30) | WNV<br>(n=13) | ND<br>(n=30) | Mean | Runtime<sup>a</sup><br>h:mm:ss<sup>a</sup> |
| --- | --- | --- | --- | --- | --- | --- | --- |
| **cytometree**<sup>1</sup> | 0.84 | 0.90 | 0.93 | 0.83 | 0.88 | 0.88 | 00:01:31<sup>†</sup> |
| **cytometree**<sup>2</sup> | 0.88 | 0.94 | 0.93 | 0.84 | 0.89 | 0.90 | 00:01:24<sup>†</sup> |
| **cytometree**<sup>3</sup> | 0.92 | 0.94 | 0.94 | 0.88 | 0.89 | 0.91 | 00:01:13<sup>†</sup> |
| **cytometree**<sup>4</sup> | 0.94 | 0.95 | 0.93 | 0.90 | 0.89 | 0.92 | 00:01:06<sup>†</sup> |
| **cytometree**<sup>5</sup> | 0.92 | 0.95 | 0.91 | 0.89 | 0.89 | 0.91 | 00:01:08<sup>†</sup> |
| ADICyt | 0.81 | 0.93 | 0.93 | 0.86 | 0.92 | 0.89 | 04:50:37 |
| flowMeans | 0.88 | 0.92 | 0.92 | 0.88 | 0.85 | 0.89 | 00:02:18 |
| FLAME | 0.85 | 0.94 | 0.91 | 0.80 | 0.90 | 0.88 | 00:04:20 |
| Flock | 0.84 | 0.86 | 0.88 | 0.83 | 0.91 | 0.86 | 00:00:20 |

<sup>a</sup> Runtime was calculated as time per sample, as displayed in Aghaeepour *et al.* (2013)
<sup>†</sup> Time was extrapolated from flowMeans runtime<sup>a</sup> which was ran together with **cytometree** on an Intel(R) i7-4770 CPU @ 3.40 GHz.

### 3.3 HIPC T-cell panel

#### 3.3.1 cytometree obtains high *F* -measures on the HIPC T-cell panel

We first compared the *F* -measures obtained by `cytometree` and flowMeans for the nine sample files (three replicates for three samples) taking as reference the manual gating of the seven centers. The results displayed in Figure 5 show that in most cases, the *F* -measures obtained by `cytometree` were better than those obtained by flowMeans.

**Figure 5:**
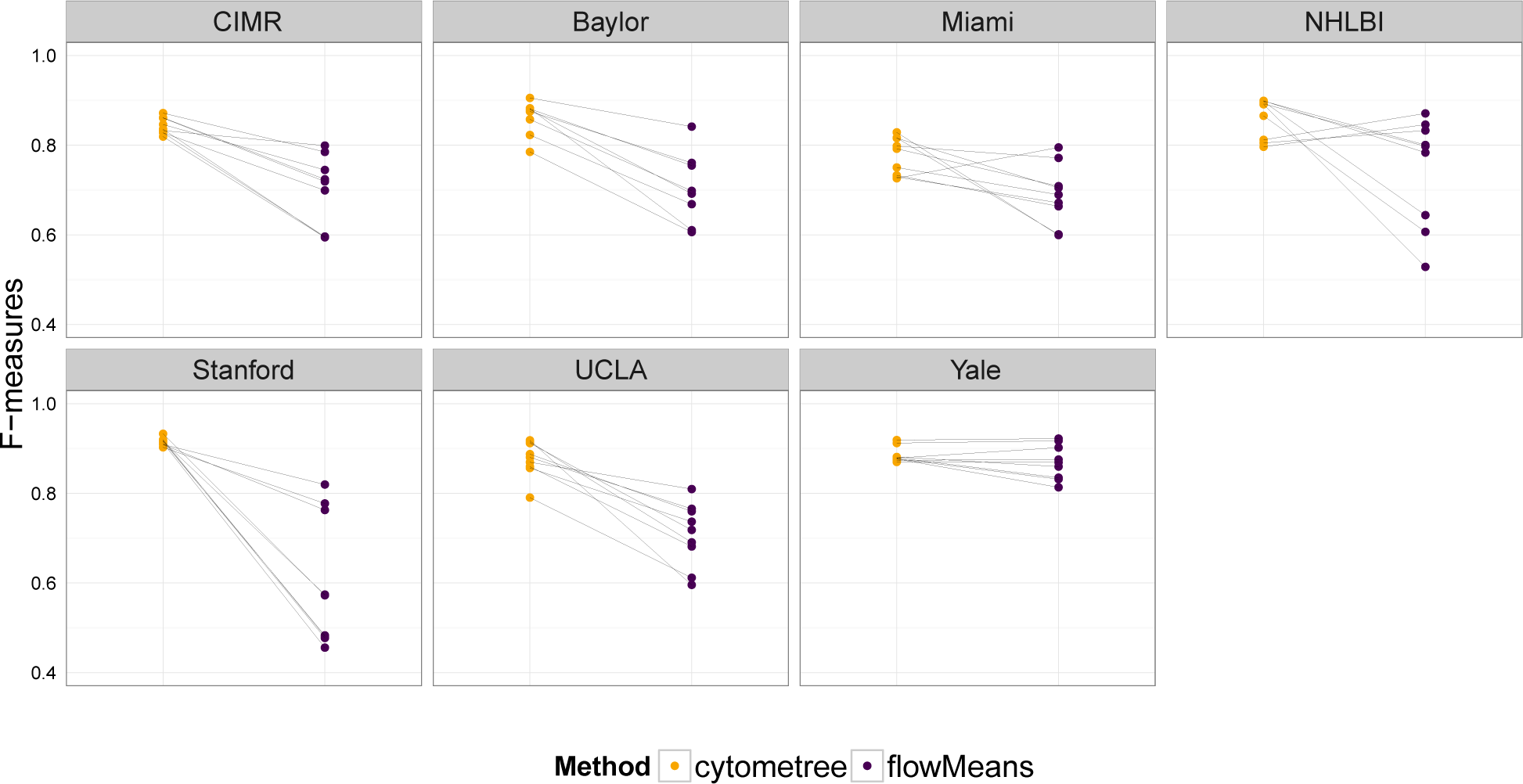
Comparison of *F* -measures, with respect to manual gating for each of the seven centers, for the nine samples of the T-cell panel of HIPC. Left and blue: cytometree; right and red: flowMeans.

#### 3.3.2 Estimation of proportions of subpopulations and their variabilities compared to manual gating

One of the goals of the method is to find proportions of pre-specified sub-populations of cells. Often the algorithm finds more subpopulations than the pre-specified ones. It is generally possible to group the finer partition that has been found to find the proportions of pre-specified populations; an example is given in Table 1. However, the algorithm has difficulties in some samples to find subpopulations representing less than 1% of the data. This is especially the case for activated T-cells, for which the number of cells can be less than 1. For this population, the variability was larger than that of the central gating.

We performed the variability analysis based on model 3. Figure 6 displays the center, biological, and residual variabilities for `cytometree` and the manual gating method for six subpopulations of the HIPC T-cell panel. The variability of `cytometree` was similar to that of manual gating, except for CD8 effector T cells. This is in line with the results presented in (Finak *et al*., 2016), where the authors showed that the CD8 effector T-cell subset was problematic due to poor separation between the HLA-DR- and HLA-DR+ populations.

**Figure 6:**
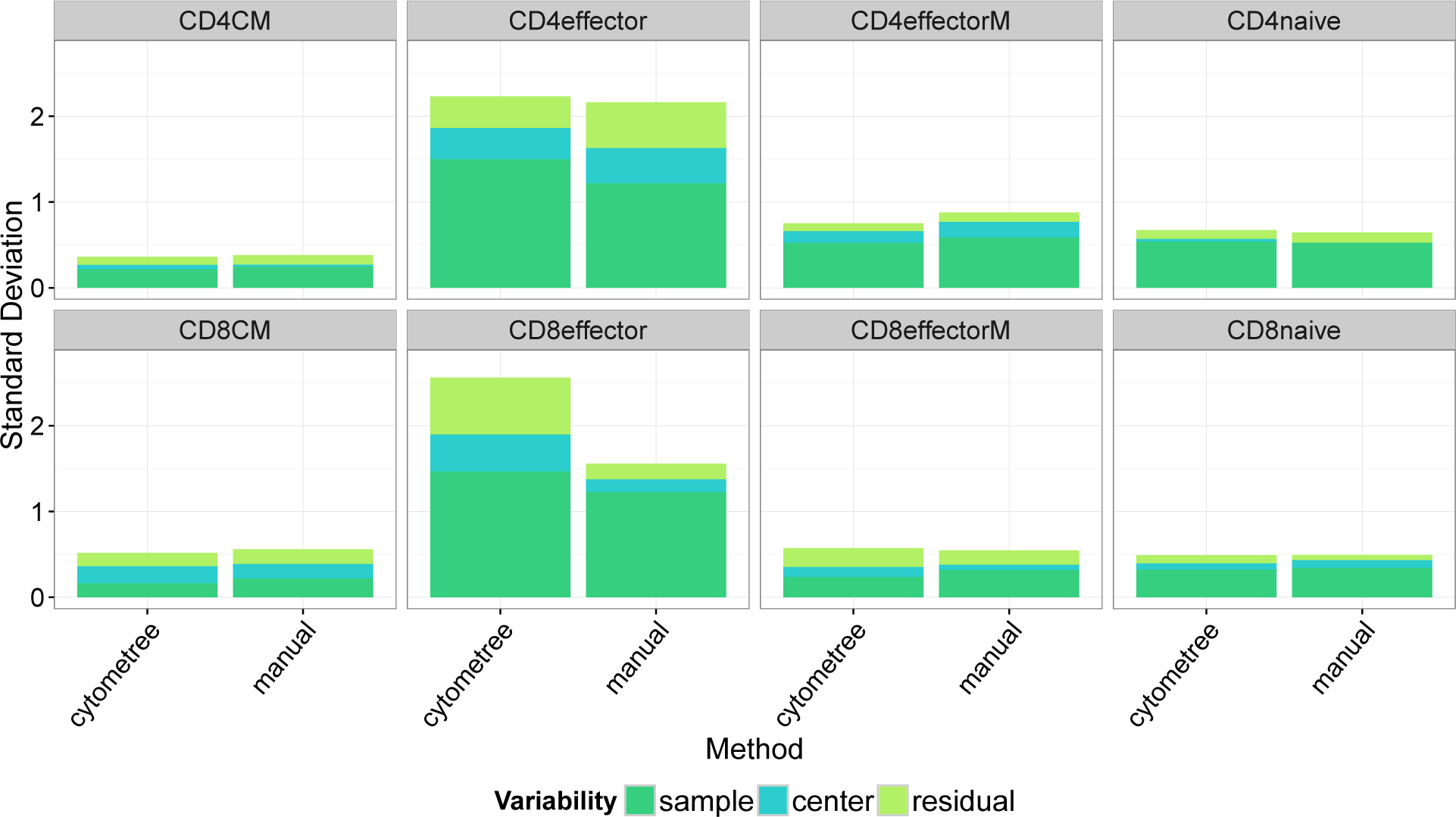
Center, biological, and residual variability for cytometree and the manual gating method for eight subpopulations of the HIPC T-cell panel.

#### 3.3.3 Discovery of new populations

As discussed previously, `cytometree` is unsupervised and defines cell sub-populations by exhaustive 1-dimensional thresholding of all markers. For example, the T-cell subpopulation labeled 9 in Figure 4 is expressing both CD4 and CD8. Although this population was likely not of primary interest in the manual analysis, `cytometree` builds sample-specific tree patterns and is able to detect such a population, which has been described in several pathological conditions as well as in normal individuals (Zuckermann, 1999; Parel and Chizzolini, 2004; Quandt *et al*., 2014).

## 4 Discussion

cytometree is an unsupervised algorithm for flow cytometry that exhibits better performance in terms of the *F* -measure than the best unsupervised algorithms, as tested on FlowCAP I data and on the HIPC T-cell panel (FlowCAP III). High *F* -measure values have been reported by Li *et al*. (2017) who proposed a deep learning algorithm, DeepCyTOF; these values, however, are not comparable to those of unsupervised algorithms; in a very recent paper, Lux *et al*. (2018) report rather modest F-measure values for DeepCyTOF. Other algorithms (Anchang *et al*., 2014; Samusik *et al*., 2016) have used binary trees, but as a secondary step; `cytometree` directly starts building the tree (see Supplemental material Sections 4 and 5).

One feature of the algorithm is its numerical simplicity and stability. In particular, mixtures of normal rather than skewed *t*-distributions (Pyne *et al*., 2009) were used; in spite of (or thanks to) this simplicity, `cytometree` obtains better *F* -measures than methods using skewed *t*-distributions for three reasons: (i) it is simpler and thus more stable; (ii) most of the distributions are not very skewed; and (iii) for moderately skewed distributions the cut-off points obtained with normal and skewed *t*-distributions are not very different. We show in Figure S2 of the Supplementary Material the results obtained with Flame and `cytometree` for one of the most skewed distribution that we have found in the diffuse large B-cell lymphoma (DLBCL) dataset; `cytometree` seems to do better than Flame on this example compared to manual gating.

cytometree is very fast and leads to population labels similar to those defined by experimentalists. This makes `cytometree` a very practical tool for experimentalists. In addition to being able to estimate proportions of pre-specified subpopulations, it can also be used in a fully unsupervised manner to perform exhaustive gating. It is fully automated and relies on a single tuning parameter, *t*^*^. We performed a sensitivity analysis to show that `cytometree` is robust to the choice of *t*^*^; the default value of 0.1 worked well in in all the samples we have tested (115 for Flowcap I and 60 from HIPC). For these reasons, `cytometree` is likely to play an important role in both clinical and discovery-based research activities.

Gating in `cytometree` is basically done through recursive thresholding of marginal densities based on the assumption that cells express or do not express certain markers, leading to bimodality. This assumption is reasonable in most scientific applications, but some markers (e.g. functional markers) might not be truly bimodal. In this case, these markers would likely not be thresholded and thus would not be represented in the gating tree. Different cases may occur, e.g. a marker may exhibit trimodality: such a feature may be retrieved through the annotation process of cytometree, as shown in Figure S1 of the Supplementary Material. A marker may be truly “continuous” and not useful for distinguishing subpopulations. Furthermore, the leaves (or any node) of the tree could then be extracted and further analyzed using other methods, including methods that have been developed to model functional markers (Lin *et al*., 2015a,b). The populations found could be further annotated using semantic labeling such as that implemented in flowCL (Courtot *et al*., 2014). Finally, it should be noted that because of the bimodality assumption `cytometree` is not adapted to gating light scatter channels (i.e. FSC and SSC) and as such it should be applied once these channels have been gated (e.g. applied to the lymphocyte population). The light scatter gates can easily be obtained by importing manual gates using the flowWorkspace package or using algorithms that have been designed to gate these two parameters (e.g. the lymphGate in the flowStats package).

As with all unsupervised algorithms, `cytometree` has difficulties in reliably identifying small populations. For instance, it correctly identified activated T cells in some samples, as shown in Table 1, but failed to identify these small populations in other samples. This result is expected as `cytometree` relies on bimodal marginal distributions to define populations. Moreover, the *D* criterion is a statistic and as such has a variance that may be large if the number of cells is small; we recommend to stop the search for population sizes lower than 50. We have done a robustness analysis showing that `cytometree` performs well in moderately small samples: results obtained on one fourth of an original sample are very similar to those obtained on the whole sample as schwn in Table 1 of Supplementary material. For rare populations, marginal density estimates are unlikely to be clearly bimodal. In such cases, some form of a priori knowledge is probably necessary. Linking the data of different samples by alignment (Cron *et al*., 2013b) or through the use of random effects as proposed by Pyne *et al*. (2014) could give more stability for rare populations.

In conclusion, the proposed algorithm is very promising in terms of its performance and its computational efficiency, both of which are important considering the pace at which the numbers of markers on single cells that can be measured is increasing.

